# Computational Design of Optimal Sequences for Targeted Hypermutagenesis Using Recombination-Coupled Diversity-Generating Retroelements

**DOI:** 10.64898/2026.06.12.731847

**Authors:** Léo Régnier, Paul Rochette, Raphaël Laurenceau, Rémi Monasson, David Bikard, Simona Cocco

**Affiliations:** CNRS UMR 8023, Laboratoire de Physique de l’École Normale Supérieure & PSL Research University, Sorbonne Université, 24 rue Lhomond, 75005 Paris, France; Institut Pasteur, Université Paris Cité, Synthetic Biology Unit, Paris, France

## Abstract

Diversity-generating retroelements (DGRs) are natural systems that accelerate evolution via targeted hypermutation at adenines. We previously developed DGRec, a system combining DGRs and recombineering for programmable mutagenesis in Escherichia coli. We here address two important issues with DGRec: the dependence of mutagenesis efficiency on the dgrRNA secondary structure and the variability of the reverse-transcription biases with sequence context and position. First, we introduce and validate a method to recode non-functional templates, i.e. with low mutagenesis efficiency, into highly functional ones through synonymous mutations. Second, we develop a Long Short-Term Memory (LSTM) model to predict DGRec mutational profiles for any given template sequence. By integrating this LSTM model with our recoding method, we establish a comprehensive workflow for customized directed evolution, enabling researchers to precisely fine-tune DGRec in vivo mutagenesis to their engineering needs.

## Introduction

Diversity-Generating Retroelements (DGRs) are specialized genomic systems that enable rapid and targeted diversification of defined loci through a reverse transcription–based mutagenic process. First discovered in the Bordetella bacteriophage BPP-1 Liu et al. [2002], DGRs have since been identified across a wide range of bacterial and archaeal genomes Coq and Ghosh [2011], Wu et al. [2017], Roux et al. [2021], Macadangdang et al. [2025], where they are thought to accelerate the evolution of phage-host recognition, as well as other bacterial functions involved in environmental adaptation.

A defining feature of diversity-generating retroelements (DGRs) is their modular organization (Fig. 1**(a)**), which consists of a template region (TR), a variable region (VR), and a dedicated reverse transcriptase (RT) Liu et al. [2002], Doulatov et al. [2004]. The diversification process initiates with the transcription of a non-coding dgrRNA. This RNA then forms a ribonucleoprotein complex with the RT and an accessory protein (Avd) Handa et al. [2025]. Within this complex, the TR serves as a stable template for RT-mediated synthesis of a mutagenized complementary DNA (TR-cDNA). The DGR RT has the unique property of making errors at a particularly high rate at adenine positions. In-vitro studies indicate that the minimal dgrRNA required for this reverse transcription includes 20 nucleotides from the avd gene, the TR, and a spacer region (Sp) Handa et al. [2018].

**Fig. 1.**
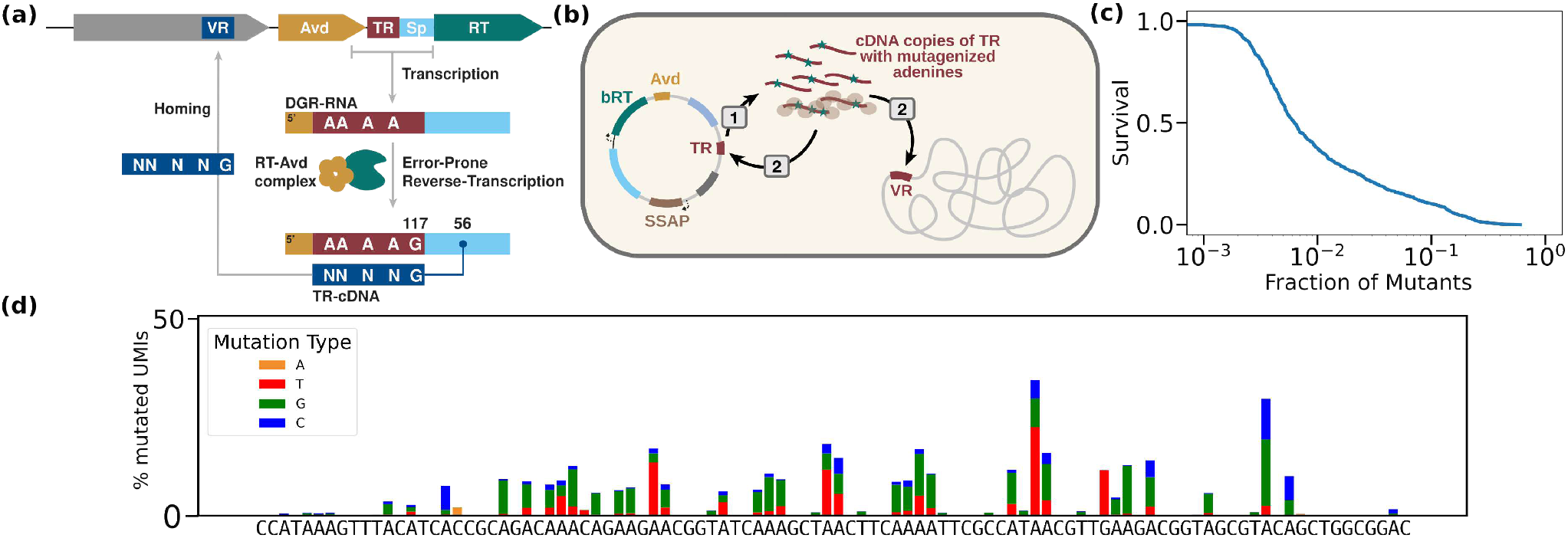
Schematic comparison of native DGR-mediated homing and DGRec-based diversification. **(a)** Native diversity-generating retroelement (DGR) architecture and homing mechanism. The nucleotides from the template region (TR) are transcribed together with nucleotides from the Spacer (Sp) and the Accessory variable determinant (Avd). The resulting dgrRNA is reverse transcribed by the reverse transcriptase (RT)-Acessory variable determinant (Avd) complex. The mutagenized TR-cDNA is subsequently integrated into the variable region (VR) through homing, replacing the VR sequence while preserving the TR. Adapted from Handa et al. [2025]. **(b)** DGRec strategy for programmable diversification. The TR can be easily replaced by Golden Gate cloning. 1) TR-cDNA are produced thanks to the DGR proteins (RT-Avd). 2) The single strand annealing protein (SSAP) CspRecT from the recombineering module, enhances the recombination of TR-cDNA to any homologous sequence, which can either be a VR in a chromosome, or the TR itself. **(c)** Survival curves i.e. fraction of sequences with percentage of mutated genotypes lower than a given threshold for the first random TRs library. **(d)** Mutational profile of a given TR obtained from the DGRec experiment.

Once synthesized, the TR-cDNA replaces the VR through an integration process, termed homing, which remains poorly understood. The VR region is typically within the C-terminal part of a protein coding gene, such as the major tropism determinant (Mtd) locus in the BPP-1 phage Liu et al. [2002]. Targeted replacement of the VR by the mutagenic cDNA drives extensive sequence diversity that ultimately modulates binding specificity and function.

We recently developped DGRec Rochette et al. [2026], a system that bypasses native homing by coupling DGR-mediated mutagenesis to recombineering (Fig. 1**(b)**). Briefly, the DGRec system consists of a plasmid comprising the Bordetella RT (bRT) and Avd coding sequences, alongside the recombineering module from the pORTMAGE system. The dgrRNA is also expressed and engineered with type IIS restriction sites, facilitating the easy modification of the TR sequence via Golden Gate cloning. Once synthesized by the bRT, the mutagenized TR-cDNA can then recombine with any homologous DNA. This process works regarless of the location of the target VR, whether on a plasmid, in the chromosome, or on a phage. Notably, unlike natural DGR systems in which retro-homing occurs at the VR but not the TR, in the DGRec method the TR-cDNA can also recombine with the TR sequence. Using DGRec, nearly any 50–200 bp sequence can be diversified.

In this paper, we address two issues raised by our previous work on DGRec. First, high-throughput DGRec experiments revealed that not all sequences function equivalently as TRs. As illustrated in Fig. 1**(c)**, in an initial library of random TRs, approximately 40% exhibit a mutation rate below 1%. This suggests that, for most TR sequences, the full DGRec process — transcription, reverse transcription, and plasmid recombination — rarely completes successfully. A likely reason identified in Rochette et al. [2026] is that the local RNA secondary structure and folding stability strongly influence the ability of the RT to read the dgrRNA, impacting overall mutagenesis efficiency. Building upon these findings, we hereafter quantitatively analyze how the energetic features of dgrRNA structure shape mutagenesis outcomes, thereby improving the accuracy of models predicting mutagenesis efficiency. Furthermore, we demonstrate that a nonfunctional TR can be recoded into an efficient one by making a few synonymous mutations, significantly expanding the scope of targetable sequences. These results deepen our understanding of the biological and technical constraints underlying DGRec design, enabling us to optimize the DGRec setup through simple modifications, such as shifting the position of the TR sequence, modifying its size or introducing synonymous mutations.

Second, beyond TR processivity, RT activity introduces intrinsic mutational biases, including context-dependent substitution patterns and mutational cascades that shape the diversity of VRs Liu et al. [2002], Handa et al. [2020], Rochette et al. [2026], Unlu et al. [2026]. The mutational profile of diversified TRs (exemple shown in Fig. 1**(d)**), reveals position and context-specific biases. We could demonstrate how the error profile of the RT (probability with which it incorporates different bases) is strongly shaped by the local sequence context. In addition, the recombineering process driving cDNA homing makes it more likely to mutate positions at the center of VR than at the edges. Our data also indicated that the DGR-RT is likely not very processive and struggles to produce long cDNA above 200 base pairs, making mutations less likely far from the RT initiation site Rochette et al. [2026]. To model these combined effects, we herefater introduce a Long Short-Term Memory (LSTM) architecture, which is particularly well-suited for capturing the sequential dependencies inherent in the DGRec system. Our model takes TR sequences as input and predicts the probabilities of generating specific VR sequences by incorporating both sequence context and RT processivity. This data-driven framework successfully captures complex positional and context dependencies in mutation profiles, providing a robust predictive model for DGRec-mediated diversification.

Combining our recoding method and VR prediction model we obtain an integrated computational pipeline implemented in the DGRec python package. This framework proposes an interface for designing and optimizing DGRec experiments, allowing users to systematically explore the mutational landscape of their target sequences and design TR sequences constraining or promoting diversification at desired positions.

## Methods: Data and Models

### Experimental data

All the experimental methods for generating the TR libraries are detailed in Supplementary Information section S1.

#### Library-1

The initial library (Library-1) comprises 702 distinct 70-nucleotide TRs with randomly selected bases.

#### Library-2

TR sequences ranging from 70 to 160 nucleotides in length are generated based on the binary classification model trained with data of Library-1, as explained in section 2.2 500 TRs attaining a prediction score > 0.7 are retained, giving rise to a library with 864 distinct TRs.

#### Library-3

TR sequences are next designed using the two classifiers described in section 2.2. We build two sub-libraries, designated 3A and 3B. Library-3A was constructed by selecting the 500 highest-scoring and 400 lowest-scoring TRs from a list of genes of interest: Cas9, iSpyMac, Lambda phages (K1F,K5, KL106, P2, Ur, V10), nanobodies (VHH 1.26, VHH 1.29, VHH 72, VHH C24, VHH F10), SlugCas9 (NNG variant), T4. To minimize sequence overlap for a given TR length, we require that both the start and end positions of each TR differ by at least 5 nucleotides. An additional 100 TRs from previous libraries are included as internal controls for cross-experiment comparison, bringing the total to 1000 TRs for Library-3A. In Library-3B, the 400 lowest-scoring TRs from Library-3A are optimized according to the methodology outlined in Section 3.1. Similarly to the first sub-library, 100 control TRs from prior experiments are incorporated, reaching a total of 500 TRs for Library-3B.

### Prediction of TR mutagenesis efficiency

In this section, we describe the method used to model the sequence dependence of the percentage of mutated genotypes in TRs, as illustrated in Fig. 1**(c)**.

We begin by analyzing Library-1, categorizing TRs based on their mutagenesis efficiency into two classes: non-mutagenic TRs (percentage of mutated genotypes < 0.3%; 100 sequences), highly mutagenic TRs (percentage of mutated genotypes > 10%; 44 sequences). Assuming that RNA structure is the primary driver of mutagenesis variability Handa et al. [2018, 2021, 2025], Rochette et al. [2026], we extract structural properties for each TR-RNA sequence using the ViennaRNA package Lorenz et al. [2011]. We then train a single-feature linear classifier based solely on the ensemble free-energy difference *ΔE*_*T R*+*Sp*_ *= E*_*T R*+*Sp*_ − *E*_*T R*_, where *E*_*T R*_ and *E*_*T R*+*Sp*_ represent the ensemble free energies of, respectively, the TR alone and the TR concatenated with the first 30 bases of the Sp. The rationale for this choice is detailed in the Supplementary Information section S2. Despite its simplicity, this model offers a key advantage: scalability to longer sequences. To optimize the Sp window, we test classifiers with Sp lengths ranging from 15 to 35 nucleotides. Performance, measured by the AUC, remain consistent across these lengths and was comparable to that of more complex models with additional features (see Supplementary Information section S2.1). Consequently, we standardize the computation of Δ*E*_*T R*+*Sp*_ using the first 30 nucleotides of the Sp.

We next apply our methodology to Library-2, a dataset enriched for TRs with high Δ*E*_*T R*+*Sp*_ values (classifier scores > 0.7). Using the same approach to determine the percentage of mutated genotypes per TR, we generate a new dataset with 460 TRs. The average mutation rate in this dataset is significantly higher than in Library-1 (12% vs. 2.5%), underscoring the importance of selecting TRs with minimal Sp interactions. TRs in Library-2 are categorized as: moderately mutagenic (label 0; percentage of mutated genotypes < 2%), highly mutagenic (label 1; percentage of mutated genotypes > 10%). To account for the remaining variability in mutation rates, we identify an additional structural feature and train a new single-feature linear classifier based on the ensemble free-energy difference Δ*E*_*Avd*+*T R*+*Sp*_ = *E*_*Avd*+*T R*+*Sp*_ − *E*_*T R*_, where *E*_*Avd*+*T R*+*Sp*_ is computed with the concatenated sequence of the Avd, TR, and the first 15 bases of the Sp. As above, larger models yielded similar performance (see Supplementary Information section S2.2).

### Prediction of TR-to-VR mutation profiles

We now aim to model the random generation of VR sequences from a given TR. Our goal is to predict the space of reachable sequences using DGRec similarly to the mutational profile in Fig. 1**(d)**. To achieve this objective, a LSTM architecture is introduced to generate VR sequences from TR inputs. LSTM are well-suited for this task: they capture long-term dependencies, are compact and efficient for small datasets, apply a convolutional structure (same kernel for each position along the sequence), adapt seamlessly to variable-length sequences, and enable rapid, flexible generation of random sequences. The two-layer LSTM model, implemented in Python, is trained on TR sequences from Library-3 with a mutation rate exceeding 3%.

#### Architecture of the two-layer LSTM model

The architecture of the LSTM used for the TR to VR generation task is the following:

- The first layer takes as input the one-hot encoded TR sequence as well as the distance (in numbers of nucleotides) of each nucleotide from the 5’ end. Because the RT proceeds from the 5’ to the 3’ end, the TR is reversed. The first layer is a bidirectional LSTM. The number of LSTM hidden-units per direction is chosen to be *k*_*BiLST M*_ = 8.
- Taking the output of the bidirectional LSTM, we apply a time distributed dense layer, outputting a vector of length *v*_*soft*_ = 6 with softmax activation and a vector of length *v*_*ReLU*_ = 2 with ReLU activation.
- We concatenate these two vectors with the one-hot encoded initial sequence. Additionally, we include the one-hot encoded TR and VR sequences shifted by one nucleotide, which are also concatenated to the previous vector. Finally, we add one dimension to mark the sequence start (initial RT position). This input to the second layer enables the model to predict future mutations based on preceding ones.
- Using the previous concatenated vector, we apply a LSTM (not bidirectional). The number of LSTM hidden-units is *k*_*LST M*_ = 16.
- As a last step, we take the output of the LSTM and apply a time distributed dense layer with softmax. The output is a vector of length 4, which is a one-hot encoding of the proposed VR nucleotide (in reversed order, as we initially reversed the TR).

The architecture of the model is represented in Fig. 2.

**Fig. 2.**
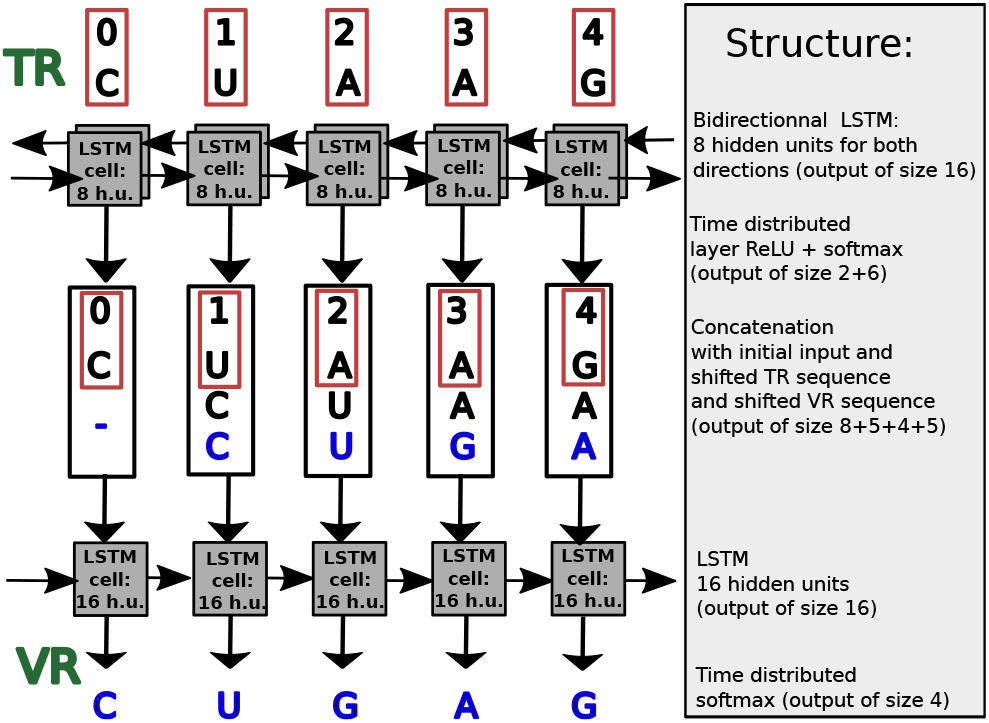
LSTM architecture. Sketch of the LSTM architecture, from top (the input) to bottom (the output). Each square represents a LSTM cell carrying out the non-linear operations. The numbers of hidden units (h.u.) of the cells are indicated in the corresponding squares.

#### VR generation

We train the TR–to–VR model on the DGRec data and test its ability to generate mutated sequences from a TR template.

To train the model, we choose TR sequences of varying lengths from Library-3, partitioning those with percentage of mutated genotypes exceeding 3% into training (80%) and test (20%) sets. For each TR, we sample 400 VR sequences from the experimental distribution to generate the (TR, VR) pairs required for training.

Then, to generate new VR from a given TR, we proceed as follows, see Fig. 2:

- First, we apply the bidirectional LSTM to the reversed TR.
- Second, we proceed sequentially, using at each step the VR nucleotide output at the previous step. We concatenate it to the previous TR nucleotide, the actual TR nucleotide (for which we want to predict the VR nucleotide) as well as the output of the bidirectional LSTM layer. We use the output vector of the second LSTM (with time distributed dense layer with both softmax and ReLU activations) to draw probabilistically the new VR nucleotide (with weight proportional to the output vector). We then move to the next TR nucleotide.

We now justify the pipeline described above. We train a model to predict, given the whole TR sequence *x* and the truncated VR sequence *y*_:*i*_ (the first *i* nucleotides in reverse transcription order), the next VR sequence *y*_*i*_. The nucleotide *y*_*i*_ is one-hot encoded, as a vector of length 4. *f*_*θ*_(*x, y*_:*i*_), the softmax output of our model of parameters *θ*, is a vector of length 4 with positive entries summing to 1. The model is trained to minimize the loss

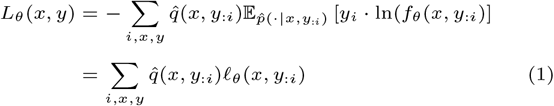

where 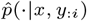 is the empirical distribution of *y*_*i*_ knowing *x* and *y*_:*i*_ (a vector with the same format as *f*_*θ*_(*x, y*_:*i*_)) and 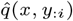 is the joint empirical distribution of (*TR, V R*_:*i*_) = (*x, y*_:*i*_) in the training set (*V R*_:*i*_ denotes the first *i* nucleotides of the VR). To minimize the loss *L*_*θ*_, one must minimize ℓ_*θ*_(*x, y*_:*i*_) for every *i*. The functional derivative with respect to *f*_*θ*_(*x, y*_:*i*_) (enforcing that the *f*_*θ*_(*x, y*_:*i*_) entries sum up to one with the Lagrange multiplier *λ*), gives

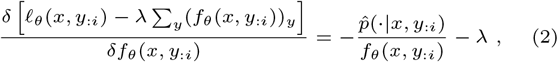

vanishes when 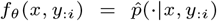 (using that both are probability distribution, hence *λ* = −1). We conclude that minimizing the total loss makes the output of our model *f*_*θ*_(*x, y*_:*i*_) minimize its distance to 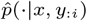.

This result can be alternatively found by rewriting the loss as

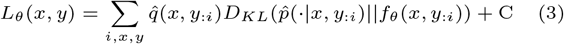

where C is independent of *θ*, and *D*_*KL*_(*p*||*f* ) is the Kullbach-Leibler divergence between the distributions *p* and *f* . Minimizing *L*_*θ*_ is the same as minimizing the divergence between distributions 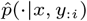 and *f*_*θ*_(*x, y*_:*i*_). 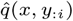 plays the role of a weight, favoring motifs (*x, y*_:*i*_) with higher occurences.

#### Results of the training

To avoid overfitting, we train different LSTM architectures changing the parameters *k*_*BiLST M*_ and *k*_*LST M*_ . The test loss are reported in Fig. 3. The best architecture corresponds to the one with *k*_*BiLST M*_ = 8 and *k*_*LST M*_ = 16 for which both the test loss and the number of parameters are minimal (for similar test loss). We use this architecture in the following.

**Fig. 3.**
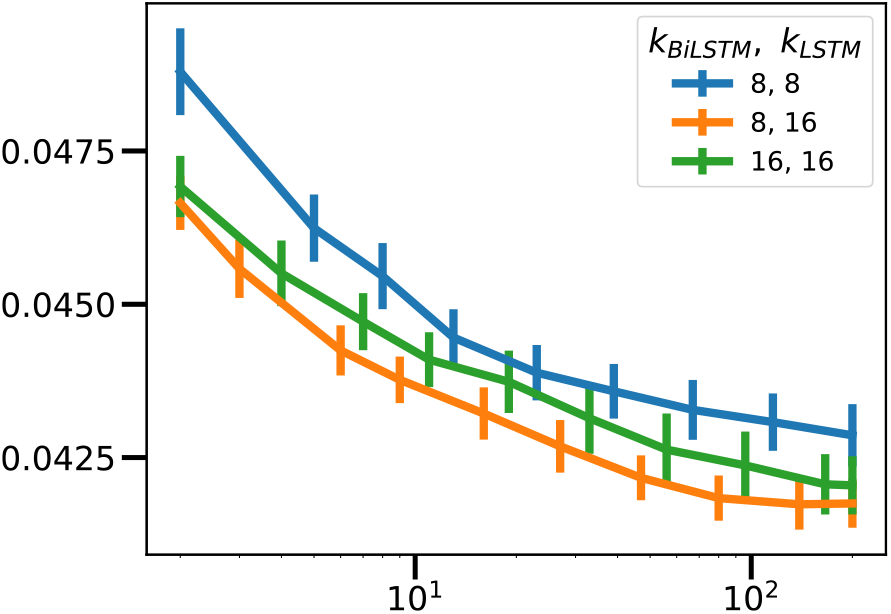
Average LSTM test loss and error bars for different LSTM architectures, changing (*k*_*BiLST M*_, *k*_*LST M*_ ). We used 20 random train-test split of the dataset for estimating the averages and the standard deviations.

## Results

In this section, we present the key results of this study. Building on our previous observation that the secondary structure of dgrRNA significantly influences DGRec mutagenesis efficiency, we first revisited the thermodynamic descriptor introduced in Rochette et al. [2026]. Consistent with our earlier findings, this feature effectively distinguishes highly mutagenic TRs from poorly mutagenic ones within Library-1. Extending this work, we identify additional thermodynamic descriptors capable of classifying TRs based on their varying percentages of mutated genotypes in the DGRec system. We further demonstrate that synonymous recoding — using the results of the combined classifiers — can optimize TR sequences to enhance their mutagenic efficiency. Finally, we develop a generative model to predict the distribution of VR sequences produced from a given TR, capturing the biases introduced by reverse transcription.

### Performances for the identifications of high mutagenic TR sequences

In sec. 2.2, we identify a key thermodynamic feature that distinguishes between TRs with high and low percentage of mutated genotypes within an initial library of random TRs (Library-1). This feature is the Gibbs ensemble free energy difference (Δ*E*_*T R*+*Sp*_ = *E*_*T R*+*Sp*_ − *E*_*T R*_) between the isolated TR sequence (*E*_*T R*_) and the TR sequence concatenated with the first 30 nucleotides of the Sp (*E*_*T R*+*Sp*_), which encompasses the template–primer region. All ensemble free energies are calculated utilizing the ViennaRNA package Lorenz et al. [2011], Hofacker et al. [1994]. As shown in Fig.4 **(a)**, the distribution of the energy-based score Δ*E*_*T R*+*Sp*_ reveals a clear distinction between TRs with low and high percentages of mutated genotypes in Library-1. Additionally, **(b)** presents the receiver-operating-characteristic area under the curve illustrating its performance with an AUC of 0.81 (95% CI: 0.68–0.91).

From a structural perspective, a large Δ*E*_*T R*+*Sp*_ indicates that the TR+Sp hybrid is less stable than the isolated TR, suggesting that the Sp nucleotides preferentially engage in self-interactions rather than base-pairing with the TR (Fig. 4**(e)**). Conversely, low Δ*E*_*T R*+*Sp*_ values denote stable TR+Sp conformations wherein the Sp is structurally sequestered by the TR (Fig. 4**(f)**). These observations support the hypothesis that efficient recruitment and processing by the RT require the TR to remain weakly coupled to its template–primer region. This hypothesis is subsequently validated using a second library (Library-2), comprising TR sequences of variable length (80–160 nucleotides) that are specifically selected for high Δ*E*_*T R*+*Sp*_ values and subjected to the same DGRec workflow as Library-1.

**Fig. 4.**
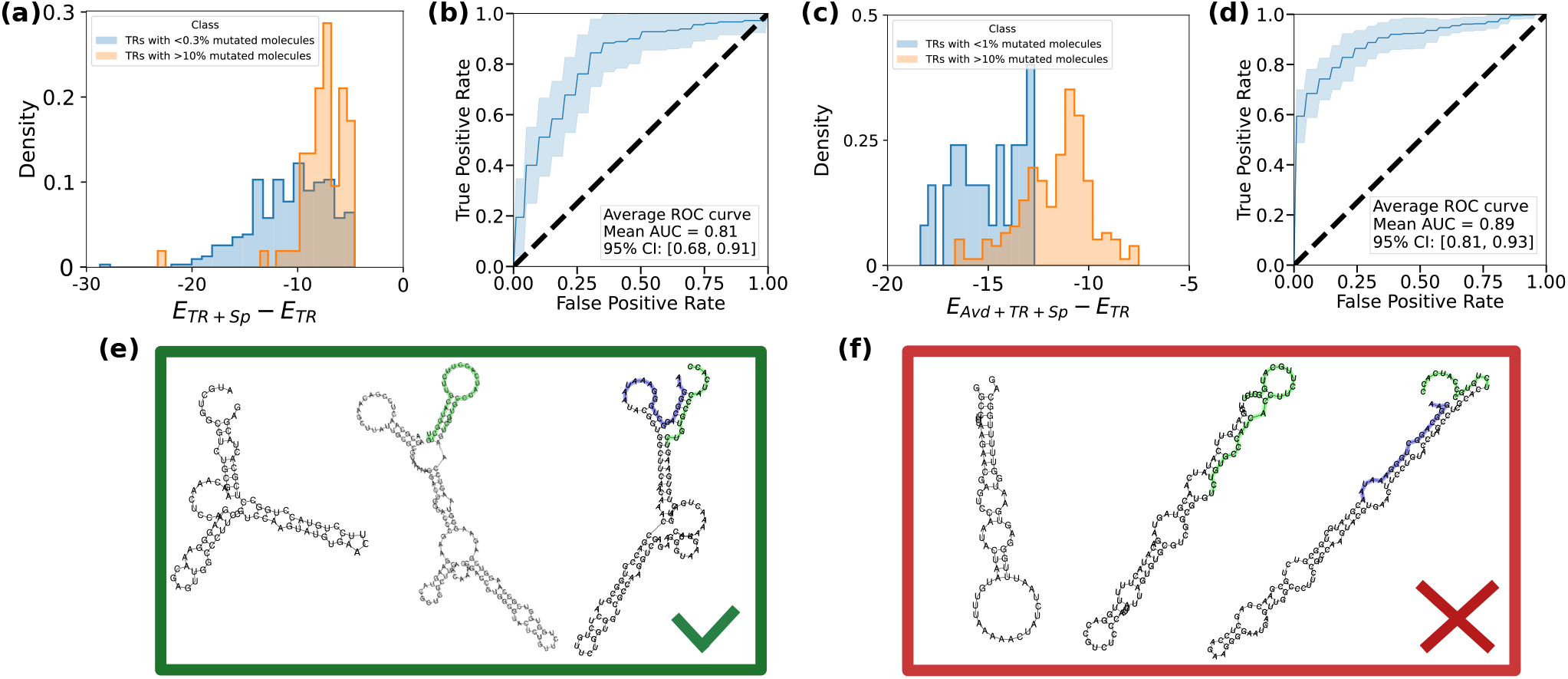
Identification of highly mutagenic TR sequences. **(a)** Distribution of the energy-based score Δ*E*_*T R*+*Sp*_ = *E*_*T R*+*Sp*_ − *E*_*T R*_ for TRs exhibiting low (< 0.3%) and high (> 10%) percentage of mutated genotypes in the initial random library (Library-1). **(b)** Classifier of Library-1 between low (< 0.3%) and high (5%) percentage of mutated genotypes TRs, representation of the receiver-operating-characteristic area under the curve. **(c)** Distribution of the combined score Δ*E*_*Avd*+*T R*+*Sp*_ = *E*_*Avd*+*T R*+*Sp*_ − *E*_*T R*_ for TRs with low (< 2%) and high (> 10%) percentage of mutated genotypes within Library-2. **(d)** Classifier of Library-2 between low (< 2%) and high (> 10%) percentage of mutated genotypes TRs, representation of the receiver-operating-characteristic area under the curve. **(e)** Representative RNA secondary structures of TRs with high percentage of mutated genotypes (TR alone, TR with 30 Sp bases in green, TR with Avd in blue and 15 Sp bases in green). **(f)** Representative RNA secondary structures of TRs with low percentage of mutated genotypes (TR alone, TR with 30 Sp bases in green, TR with Avd in blue and 15 Sp bases in green).

Library-2 comprises 460 TR sequences predicted by the classifier to have high percentage of mutated genotypes (prediction score > 0.7). It yielded 144 highly mutagenic TRs (percentage of mutated genotypes > 10%), which represents 31% of the library, i.e. a four-fold gain relative to Library-1. Notably, non-mutagenic TRs (percentage of mutated genotypes < 0.3%) are not detected in Library-2. Nevertheless, a substantial subset of TRs exhibited only moderate mutagenic activity, with fewer than 5% mutated genotypes, highlighting significant potential for further optimization. Consequently, our subsequent objective is to discriminate between moderately and highly mutagenic TRs within the Library-2 dataset.

As anticipated, the original descriptor Δ*E*_*T R*+*Sp*_ lacks robust predictive power in this refined dataset. A subsequent round of feature selection and cross-validation identifies an additional thermodynamic descriptor, Δ*E*_*Avd*+*T R*+*Sp*_ = *E*_*Avd*+*T R*+*Sp*_ −*E*_*T R*_, where *E*_*Avd*+*T R*+*Sp*_ denotes the ensemble free energy of the concatenated sequence comprising the 20-nucleotide Avd region, the TR, and the initial 15 nucleotides of the Sp. As illustrated in Fig. 4**c**, Δ*E*_*Avd*+*T R*+*Sp*_ successfully discriminates between moderately and highly mutagenic TRs within Library-2. Furthermore, the classifier employing Δ*E*_*Avd*+*T R*+*Sp*_ as the sole predictive feature achieves an AUC of 0.89 (95% CI: 0.81–0.93) as shown in Fig 4**(d)**. From a structural perspective, high values of Δ*E*_*Avd*+*T R*+*Sp*_ correspond to conformations wherein the Avd-RNA and Sp-RNA remain accessible (see Fig. 4**(e)**), facilitating mutual interaction. Conversely, low values suggest that both regions are structurally sequestered via base-pairing with the TR (see Fig 4**(f)**). This interpretation aligns with the results derived from Library-1 and corroborates previous studies demonstrating that efficient mutagenic reverse transcription requires Sp–Sp and Avd–Sp interactions Handa et al. [2025].

Altogether, our results suggest that when either the Sp or Avd region is structurally sequestered via base-pairing with the TR (indicated by low Δ*E*_*T R*+*Sp*_ or low Δ*E*_*Avd*+*T R*+*Sp*_ values), mutagenic reverse transcription activity is markedly reduced. Conversely, RNAs exhibiting high values for both descriptors adopt conformations that promote Sp–Sp or Avd–Sp interactions, thereby facilitating productive reverse transcription.

In summary, optimal predictive performance is achieved by employing a dgrRNA model that integrates both Δ*E*_*T R*+*Sp*_ and Δ*E*_*Avd*+*T R*+*Sp*_, as these descriptors capture complementary aspects of the thermodynamic landscape governing TR efficacy.

### Synonymous recoding of TR sequences offers highly mutagenic TR

We subsequently develop a third library (Library-3) by jointly applying both thermodynamic descriptors, selecting sequences that exhibit sufficiently high values for both Δ*E*_*T R*+*Sp*_ and Δ*E*_*Avd*+*T R*+*Sp*_. As depicted in Figure 5, this finalized library shows a marked increase in overall fraction of mutated genotypes; more than half of the TRs were highly active (over 10% mutated genotypes).

**Fig. 5.**
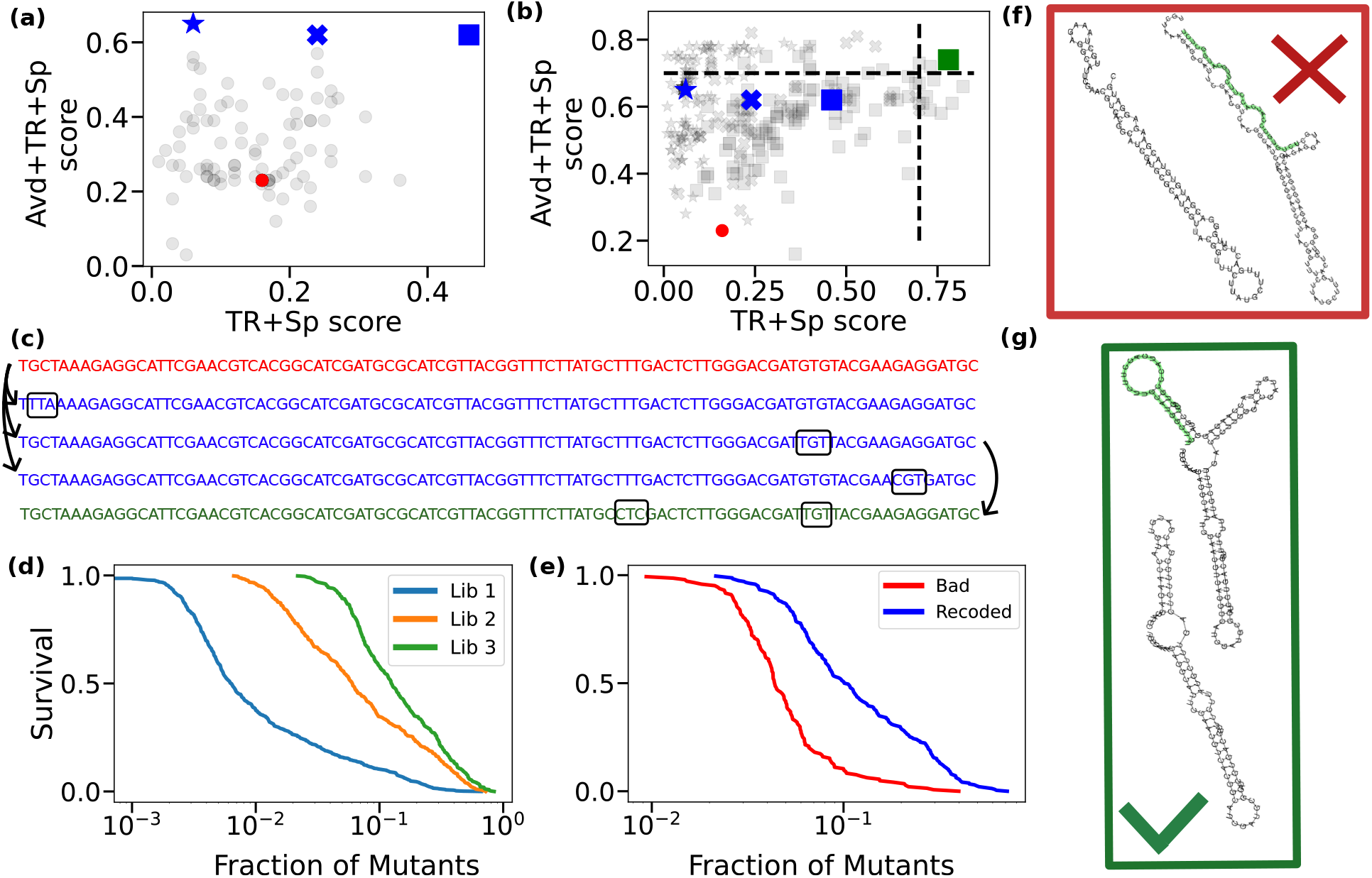
Results of the successive TR optimizations. **(a)** and **(b)** Scatter plots of the Δ*E*_*T R*+*Sp*_ and Δ*E*_*Avd*+*T R*+*Sp*_ scores for variants of the initially low score TR. **(a)** Starting from the initial TR with low scores (red circle), we estimate the scores of all one synonymous codon change variants (black circles) and identify the Pareto-optimal ones (blue cross, square and star), for which no other variant has both higher Δ*E*_*T R*+*Sp*_ and Δ*E*_*Avd*+*T R*+*Sp*_ scores. Since none of these Pareto-optimal variants achieve scores greater than 0.7, the procedure continues. **(b)** By computing the scores of all one-codon change variants derived from the previous Pareto-optimal sequences (in black, with shapes matching the optimum it comes from), we identify a variant (in green square) with both scores exceeding 0.7. This variant is the final recoded TR sequence. The nucleotide sequences corresponding to the initial TR, Pareto-optimal variants, and the final recoded sequence are shown below the plots, with their codon replacements framed. **(c)** Successive sequences of interest of the optimization: in red the initial sequence, in blue the three starting one-codon change variants for the second step, in green the final sequence. **(d)** Survival curves i.e. fraction of sequences with fraction of mutated genotypes lower than a given threshold for Libraries 1–3. **(e)** Survival curves comparing TRs predicted to have low fraction of mutated genotypes before (Bad) and after synonymous recoding (Recoded). **(f)** Secondary structure of a TR with low fraction of mutated genotypes. Both the structures with or without the 30 first nucleotides of the Sp are shown. **(g)** Secondary structure of a TR with initially low fraction of mutated genotypes recoded to have a high fraction of mutated genotypes. Both the structures with or without the 30 first nucleotides of the Sp are shown. In both **(f)** and **(g)** the Sp sequence is highlighted in green.

We then investigated whether poor TR sequences can be rationally recoded to achieve high mutagenesis rates while keeping the amino acid sequence fixed, in order to preserve the VR-encoded protein that the TR will produce via DGRec.We devised an optimization strategy based on iterative synonymous recoding. Starting from a TR sequence that matches a natural coding sequence, we systematically evaluate all potential variants containing a single synonymous codon substitution. If no single variant achieves a prediction score above 0.7 for both the Δ*E*_*T R*+*Sp*_ and Δ*E*_*Avd*+*T R*+*Sp*_ classifiers, we select the optimal variants—defined as those for which no other variant exhibits superior performance across both metrics. This process is repeated iteratively, permitting up to six synonymous codon substitutions relative to the original sequence. In instances where no variant satisfies the 0.7 threshold for both classifiers, we retain the sequence that maximizes Δ*E*_*T R*+*Sp*_. As outlined in Figure 5, this iterative recoding provides a highly effective framework for enhancing TR processing by the DGRec system. Ultimately, this rational design approach yields a substantial amplification of mutagenic efficiency, achieving an average relative increase of 322% in the percentage of mutated genotypes of the recoded sequences.

Of note, in cases where the TR does not self-mutagenize but only undergoes VR replacements (as described in Sec. 1 and in Rochette et al. [2026]), optimizing a TR sequence via synonymous recoding may require compensatory adjustments to the VR sequence to preserve homology. While synonymous recoding carries minimal risk of compromising function, extensive codon substitutions in the TR can reduce recombination efficiency if the VR is not similarly adjusted, as this process critically depends on TR–VR homology. Thus, recoding the TR alone can enhance cDNA production through increased mutagenesis but may concurrently diminish recombination if the VR remains unmodified. Another strategy would be to adjust the TR window by shortening or adding nucleotides while preserving functional sequences and the region of diversification of interest.

Ultimately, this study demonstrates that TR sequences can be rationally optimized to drive robust mutagenesis within the DGR framework. As summarized in Fig 4**(d)** and 5 **(f)**, an efficient TR is defined by three principal thermodynamic and structural characteristics: *(i)* high intrinsic TR stability, *(ii)* weak interactions with Sp, and *(iii)* weak interactions with the Avd region.

### TR to VR mutation profile prediction

In the previous section, we focused on predicting and improving the mutagenesis rate of the DGRec method. Here we focus on predicting the *distribution* of mutated sequences generated by the DGRec process: given an initial TR, which VR sequences can be accessed through the reverse transcription process. This amounts to determining the conditional probability *p*(VR | TR), i.e. the probability of generating a specific VR sequence given an initial TR.

Figure 6 illustrates the ability of the LSTM model to faithfully reproduce the key statistical features observed in the experimental data and noted by Refs. Rochette et al. [2026], Unlu et al. [2026]. Panels **(a)** and **(b)** demonstrate that the model captures the strong context dependence of mutation probabilities, extending beyond nearest-neighbor effects. Panels **(c)** and **(d)** show that the model correctly reproduces mutational cascades, with elevated mutation probabilities at downstream positions following an initial mutation. Panel **(e)** demonstrates that the LSTM model accurately captures the positional dependence of mutational frequency. Finally, the nucleotide substitution biases learned by the model Panels **(f)** are consistent with those measured experimentally (Fig. 1**(d)**).

**Fig. 6.**
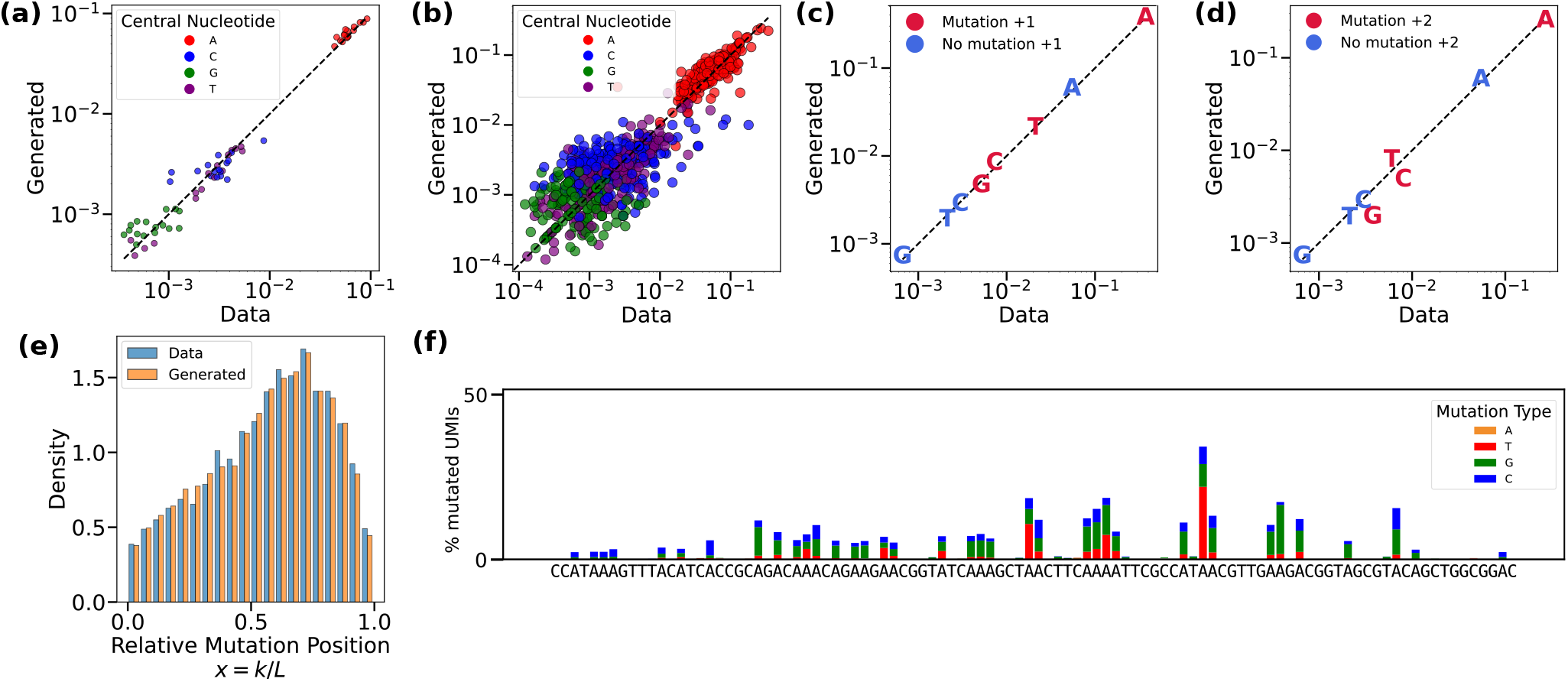
**(a)** Mutation probability of a central nucleotide as a function of its two nearest neighbors. **(b)** Mutation probability of the central nucleotide as a function of its five nearest neighbors. **(c)** Mutation probability of the immediately following nucleotide conditioned on whether the previous nucleotide has mutated. **(d)** Mutation probability at distance two conditioned on a mutation at the initial position. **(e)** Mutation profile as a function of the relative mutation position *x* = *k/L* with *L* the sequence length and *k* the mutation position. **(f)** Mutation profile of a sequence of length 100 from Library-2 generated by the LSTM model. The sequence is the same as in Fig. 1 **(d)**.

The model demonstrates strong agreement with experimental data across all tested conditions, accurately reproducing both the spatial distribution of mutations and the relative frequencies of nucleotide substitutions. Collectively, these results establish that the LSTM-based generative model effectively predicts the distribution of VR sequences accessible from a given TR. This capability enables quantitative exploration of the mutational landscape shaped by reverse transcription, providing a robust framework for further investigation.

Our LSTM-based model provides a probabilistic representation of the mutagenesis process itself, enabling the precise prediction of the VR sequence distributions generated from a given TR. Rather than relying exclusively on aggregate percentage of mutated genotypes, this approach empowers experimentalists to anticipate specific mutation profiles, nucleotide substitution biases, and correlated mutation events across the sequence. Researchers can design experiments that not only maximize overall diversification but also rationally steer the accessible sequence space toward targeted phenotypic regions. To illustrate this utility, we evaluated the amino acid frequencies generated from a TR corresponding to a sub-region of the nanobody VHH 1.29 CDR3. Targeted diversification at nanobody position 106 (corresponding to index 10 in Figure 7, a residue known to interact with the antigen) was achieved by substituting the initial TR codon with AAC. Consistent with the DGRec codon-design principles established in Ref. Rochette et al. [2026], the LSTM diversification help assess the addition of the high diversity TR triplet AAC at a strategic position.

**Fig. 7.**
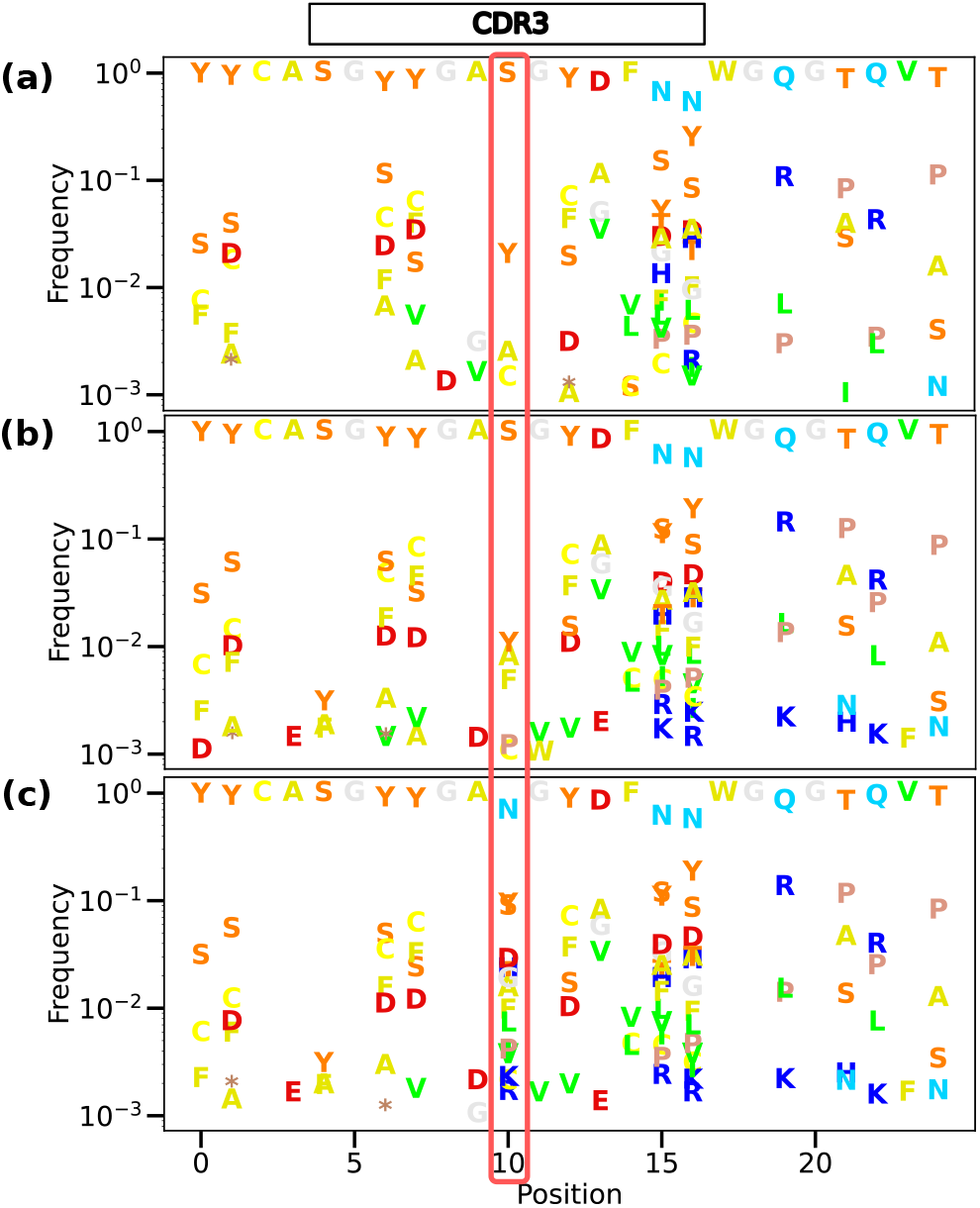
**(a)** Experimental amino acid frequencies obtained by DGRec diversification of a given T R. **( b)** Amino acid frequencies obtained by DGRec diversification of the same TR via generation of 4 ·10^4^ V Rs using the LSTM. **(c)** Amino acid frequencies obtained by DGRec diversification of the same TR but where the codon corresponding to position 10 has been changed to AAC. The 4 · 10^4^ VRs have been generated using the LSTM.

## Conclusion

In this study, we establish a predictive framework for the rational design of DGR-based diversification experiments. Leveraging the DGRec system allows diversification to be targeted to arbitrary genomic region of 50-200nt, and within these region the addition or deletion of adenines in the TR sequence can be used to control the specific residues undergoing diversification. Our analysis identifies key sequence-based criteria that facilitate the reliable selection of TRs driving a high mutagenesis rate. These predictors offer practical guidelines for constructing TR optimized for maximum diversification efficiency.

However, some amino acids are encoded by codons that can mutate into stop codons via a single adenine substitution (e.g., lysine codons AAA and AAG can become TAA and TAG). While avoiding such mutations is desirable to preserve protein functionality, this constraint may limit the optimization process. However, for lysine, all its codons can potentially become stop codons through adenine mutagenesis, making such avoidance impossible. Consequently, selecting an optimal strategy necessitates navigating inherent trade-offs among mutagenesis efficiency, recombination rates, and the functional preservation of the target protein.

Our framework, as illustrated in Figure 8, provides a structured methodology for designing TR sequences that achieve both high percentage of mutated genotypes and targeted diversification at designated positions. These computational methods have been implemented in a Python package, DGRec. By enabling the systematic, model-driven design of DGR-based diversification experiments, our framework not only facilitates the quantitative exploration of protein sequence space but also enhances the interpretability and scalability of directed evolution workflows.

**Fig. 8.**
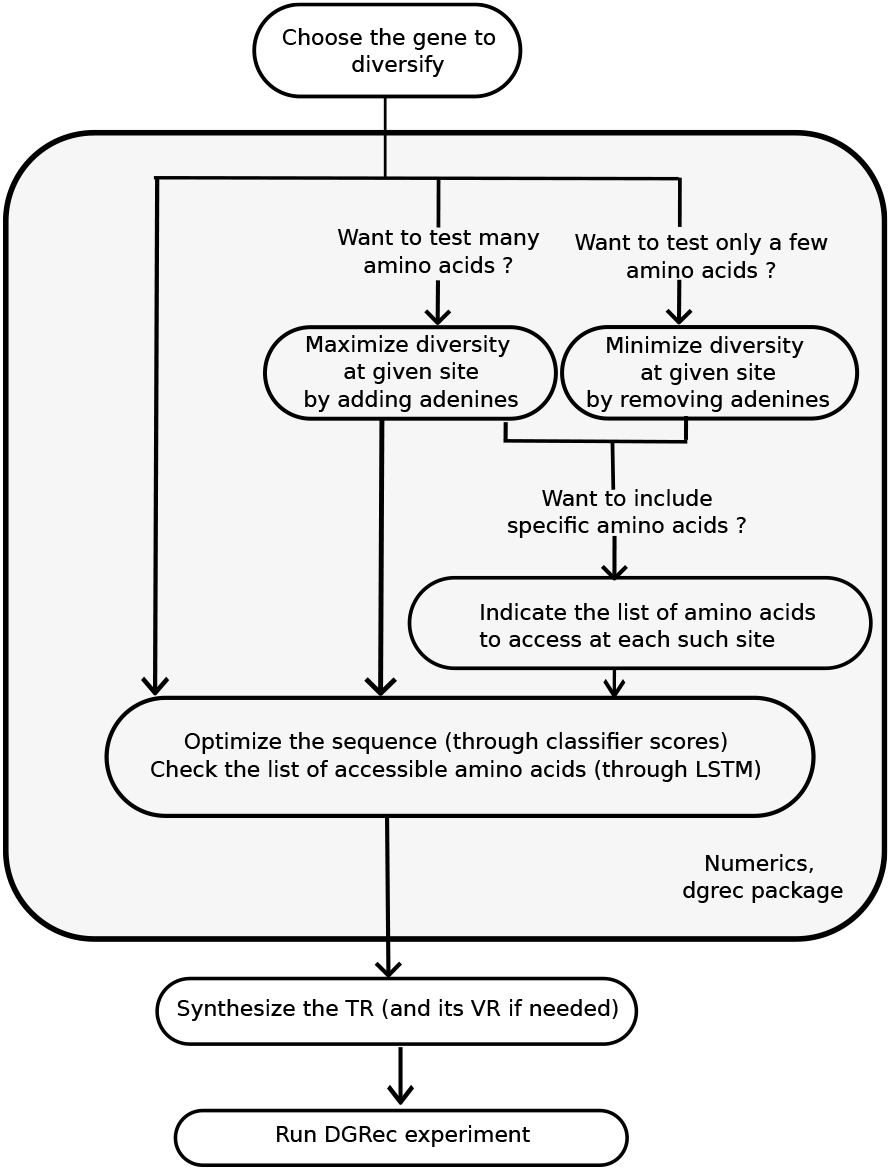
Framework for designing TR sequences that balance energy constraints and targeted diversification. The grey box highlights the informatics pipeline implemented in the DGRec package.

A notable limitation of this study, however, is its reliance on the self-mutagenesis DGRec dataset, which inherently restricts its current scope to self-mutagenesis paradigms wherein the TR and VR sequences are initially identical. Consequently, the framework does not yet account for TR–VR spatial dissociation or the impact of non-homology between these regions. Both factors may significantly influence recombination efficiency and mutational patterns in more complex, bipartite architectures.

Despite these constraints, the framework remains highly adaptable. Although currently trained exclusively on DGRec data, its underlying principles are broadly generalizable and can be extended to other DGR systems, provided sufficient sequencing data is available. Furthermore, our approach could be integrated with protein modeling tools to evaluate the functional and structural properties of DGRec variants generated in silico by the LSTM model. This would allow balancing the diversity of generated VR sequences with the functional integrity of the resulting variants. This adaptability positions our framework as a versatile tool for enhancing the precision and efficacy of protein engineering and directed evolution strategies.

In summary, this integrated framework represents a powerful tool for guiding DGRec-based experiments, bridging the gap between computational predictions and experimental outcomes. It empowers researchers to harness the full potential of DGRec-mediated diversification while maintaining control over the mutational process, ultimately facilitating more efficient and targeted evolution of biological sequences.

## Code availability

The user-friendly package to predict TR scores and to generate mutational profiles using the LSTM model is publicly available at https://dbikard.github.io/dgrec/. The repository provides implementations of the scoring pipeline, the LSTM-based prediction of mutation probabilities, and example workflows for reproducing the analyses presented in this study. The additional repository https://github.com/LeoReg/DGRec_numerics.git contains the data and the code used to train the classifiers and the LSTM model as well as for making the figures of the present work.

## Acknowledgements

We thank Ivan Lecce and Elena Lopez-Rodriguez for constructive discussions. This work was supported by the French National Research Agency (ANR) under Grant No. ANR-23-CE45-0034 and ANR-10-LABX-62-IBEID and the European Research Council (101044479).

