## Supplementary Information for "Computational Design of Optimal Sequences for Targeted Hypermutagenesis Using Recombination-Coupled Diversity-Generating Retroelements"

### CONTENTS

S1. Experimental methods 1
1. Bacterial strains, plasmids, media and growth conditions 1
2. Preparation of the TR Library-1 1
3. Preparation of the TR Library-2 2
4. Preparation of the TR Library-3 2

S2. Identifying predictive features for highly mutagenic sequences using linear classifiers 2
1. Classifier using TR Library-1 2
2. Classifier using TR Library-2 3

Supplementary References 4

### S1. EXPERIMENTAL METHODS

#### *1. Bacterial strains, plasmids, media and growth conditions*

All bacterial strains and plasmids used in this work are from Ref. [S1]. All plasmids were propagated in the the E. coli strain MG1655ΔfhuA, except for plasmids containing the ccdB toxin which were propagated in One Shot™ ccdB Survival™ 2 T1R (ThermoFisher). All the strains were grown in lysogeny broth (LB) at 37°C and shaking at 180 RPM. For solid medium, 1.5 % (w/v) agar was added to LB. The following antibiotics were added to the medium when needed: $50\mu g/ml$ kanamycin (Kan).

#### *2. Preparation of the TR Library-1*

A single-stranded oligonucleotide with two overhangs containing BsaI restriction sites and 70 degenerate bases (N) was ordered to IDT (PR123, see Supplementary Table 1 for full oligonucleotide sequences). A PCR with 5 amplification cycles was carried out to make dsDNA with PR002 and PR003 oligonucleotides. This TR was then cloned into pPR150 and transformed into E. coli MG1655 ΔSbcBΔRecJ (sRL001). Cells plated on the corresponding selective media (Kan). After an overnight growth, 768 individual colonies were picked and transferred into 1 mL of LB-Kan in a 96-well plate and allowed to grow for 6-8 hours. These un-induced pre-cultures were diluted 1000-fold into 1mL of LB with Kan and 50 $\mu M$ DAPG (inducing the bRT module) in a 96 deep-well plate, and allowed to grow for 24 hours at 37°C with shaking at 180 rpm, reaching stationary phase.This 1000-fold dilution and growth was repeated once more for all cultures to reach 48h of DGRec induction. Each 96 well plates were pooled together and plasmid DNA was extracted for each plate. Plasmid DNA was then prepared for sequencing. A PCR-1 to add the UMIs was carried out with 2 cycles of amplification with Q5polymerase (New England Biolabs) with PR121 and PR133 oligonucleotides. PCR-1 samples were treated with 1µL of ExoI (New England Biolabs) to remove single-stranded DNA leftovers. The samples were then purified with AMPure XP magnetic beads (Beckman Coulter). A barcoding PCR-2 for Illumina library preparation was performed with 15 cycles. Barcoded amplicons were then purified with AMPure XP magnetic beads (Beckman Coulter), pooled, and the final pooled library was quantified with the NEBNext Library Quant Kit for Illumina (New England Biolabs). The pooled library was mixed to a 2:1 molar ratio with PhiX v3 control (Illumina) to ensure base diversity. The samples were then sequenced on a NextSeq 1000/2000 P1 300 cycles (Illumina) with 151 cycles of paired-end sequencing covering the TR both in forward and reverse direction. In order to determine the non-mutated sequences of the TRs, we sequenced the samples before induction with DAPG, grouped the sequences by the first 7 and last 7 bases and considered as non mutated TRs the sequences where we retrieved more than 100 molecules. In a similar way, we grouped the sequences of mutated samples (after 48h induction) by the first 7 and last 7 bases and aligned them with their corresponding non mutated samples to determine the mutated genotypes. Among the 768 picked clones, we recovered 702 different TRs with more than 1000 different molecules each.

Supplementary Table 1. Oligonucleotide sequences

| **Name** | Sequence |
|---|---|
| PR002 | GGCACGTACGGTCTCGATAA |
| PR003 | GAAGATCGCGGTCTCTCAGA |
| PR121 | GTGACTGGAGTTCAGACGTGTGCTCTTCCGATCTNNNNNNNNNNCCATGCAAGAAGGTGATGGGCA |
| PR123 | GGCACGTACGGTCTCGATAAANNNNNNNNNNNNNNNNNNNNNNNNNNNNNNNNNNNNNNNNNNNNNNNNNNNTCTGAGAGACCGCGATCTTC |
| PR133 | TTCCCTACACGACGCTCTTCCGATCTNNNNNNNNNNNAAGGGCAGGCTGGGAAAT |

#### *3. Preparation of the TR Library-2*

To the TR sequences selected for Library-2, the same overhangs as in Library-1 (containing BsaI restriction sites) were added and the 500 sequences were then synthesized as a pooled oligonucleotide library by Twist Bioscience. All downstream experimental steps were performed as described for Library-1, with 864 individual colonies picked.

#### *4. Preparation of the TR Library-3*

Library-3A and Library-3B were synthesized as two separate pooled library by Twist Bioscience and all downstream experimental steps were performed as described for Library-1 with 864 individual colonies picked for each sub-library.

### S2. IDENTIFYING PREDICTIVE FEATURES FOR HIGHLY MUTAGENIC SEQUENCES USING LINEAR CLASSIFIERS

In this section, we describe the methods used to classify TR sequences based on their percentage of mutated genotypes in the DGRec setup. Using linear classifiers, we identify features that are both predictive of the DGRec mutagenic rate and biologically relevant, distinguishing sequences with high mutation rates from those with low activity. Notably, we find that folding energies emerge as the most predictive biological factors.

#### *1. Classifier using TR Library-1*

To identify the primary features governing the reverse transcription of a TR and the subsequent recombination of its cDNA with the target region via DGRec, we evaluated various feature sets and classifier architectures to explain the observed variance in mutagenesis efficiency displayed in Fig.**??** **(c)**. Sequences from Library-1 were first categorized based on their mutagenesis efficiency: non-mutagenic (TRs with $< 0.3\%$ mutated sequences; nTRs = 100) and highly mutagenic (TRs with $> 10\%$ mutated sequences; nTRs = 44). Given the limited dataset, we employed small, simple architectures to mitigate the risk of overfitting. Hypothesizing that RNA structure is the primary determinant of mutagenesis variability [S2–S4], we utilized the ViennaRNA package to extract the structural properties of each TR-RNA sequence. Specifically, we computed the ensemble free energy (`E`), minimum free energy (`MFE`), mean expected accuracy (`MEA`), positional entropy (`entropy_x`, where x is the relative position in the DGR-RNA), and maximum base-pairing probability (`max_bpp_x` with `x` the relative position in the DGR-RNA). Additionally, we calculated the differences between these thermodynamic metrics and those of the baseline dgrRNA alone, denoting these derived features with a Δ prefix. Finally, all features were extracted using windows of 70 and 100 nucleotides along dgrRNA sequence (taking steps of 20 nucleotides), window position indicated by the suffix `from_x_to_y` where `x` is the sequence start and `y` its end. This yields a comprehensive set of 3,126 features per sequence.

Using a logistic regression model with $L_1$ regularization (regularization strength = 0.05, balanced class weights), we achieved an average balanced AUC of 0.76 across 100 independent runs, utilizing a randomized 80/20 train-test split for each iteration. Energy-related features emerged as the most predictive, particularly the ensemble free energy difference, $\Delta E_{TR+Sp} = E_{TR+Sp} - E_{TR}$ (Fig. Fig. S1). This energy difference term consistently yielded the highest regression coefficient and was the sole feature invariably selected across all random training permutations.

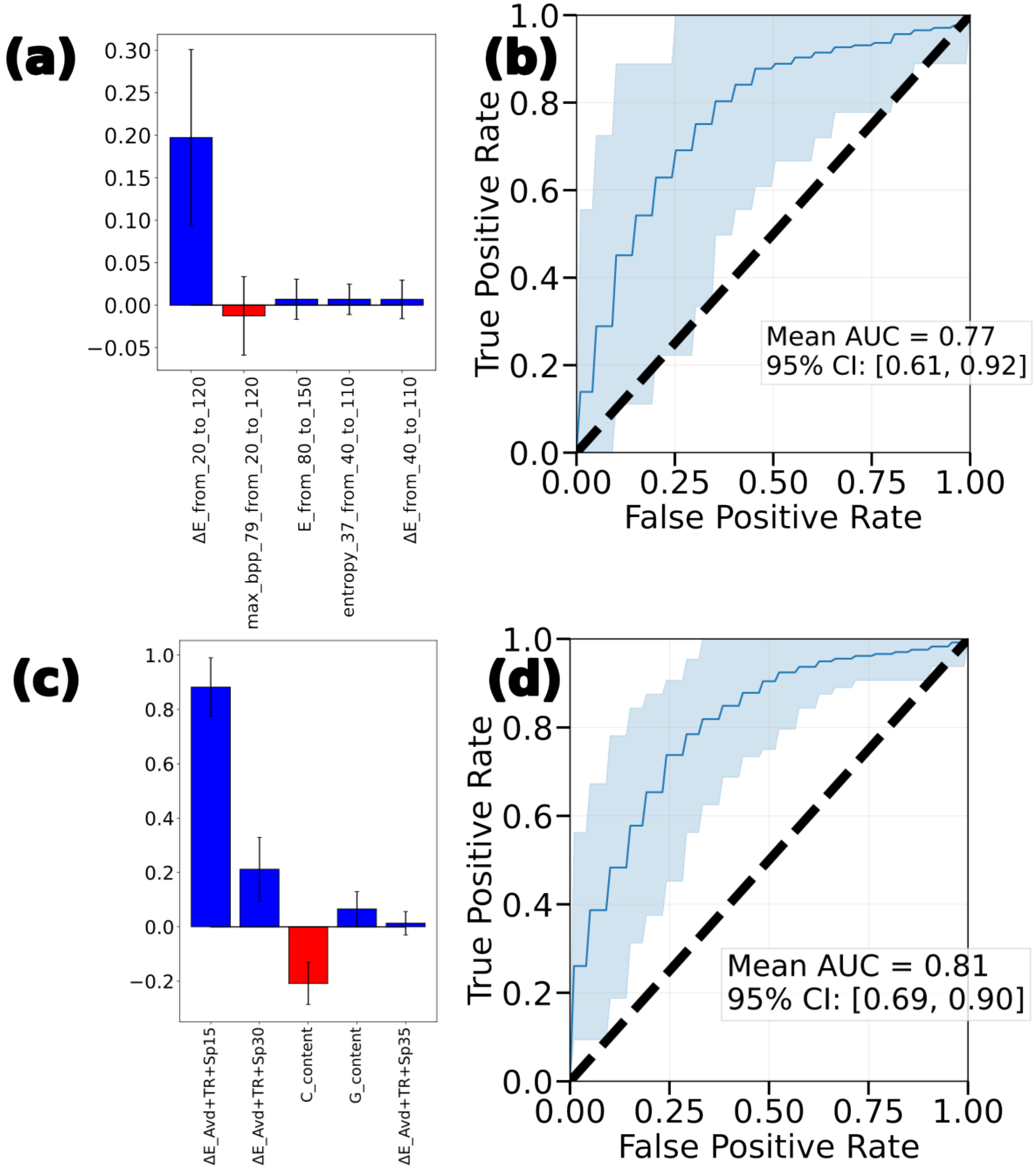


Fig. S1. **(a)** Coefficients of the $L_1$ linear regression curve sorted according to their amplitude with their standard deviation for classification of Library-1. Positive coefficients are in blue, negative in red. **(b)** Average receiver-operating-characteristic curve with its 95% interval of fluctuation for classification of Library-1. **(c)** Coefficients of the $L_1$ linear regression curve sorted according to their amplitude with their standard deviation for classification of Library-2. Positive coefficients are in blue, negative in red. **(d)** Average receiver-operating-characteristic curve with its 95% interval of fluctuation for classification of Library-2.

#### 2. *Classifier using TR Library-2*

We evaluated a novel set of length-independent features to capture characteristics that explain the differences in mutagenesis:

- **Nucleotide composition:** Overall GC content (`GC_content`), as well as individual relative frequencies of G and C nucleotides (G_content, C_content).

- **Ensemble free energy differences:** Computed between the TR alone and constructs appended with Sp windows of 10, 15, 20, 25, 30, and 35 nucleotides, with and without the 5' Avd sequence. They are denoted with the prefix ΔE and the sequences parts included in the computation as suffix.
- **TR-TR interactions:** Features derived from base-pairing probability (BPP) matrices within the TR, including the ten highest TR-TR base-pair probabilities, the ten highest marginal base-pairing probabilities per base, and the total internal base-pairing probability.
- **TR-Avd or TR-Sp interactions:** The ten highest probabilities of TR nucleotides pairing with external nucleotides from Avd or Sp (Top_TR_Other_x, where x is the rank), along with the total base-pairing probability between the TR and non-TR regions.

The results of this classification task (using an $L_1$ penalty coefficient of 0.05) are presented in Fig. Fig. S1 **(c)** and **(d)**. We identified the ensemble free energy difference involving the Avd, the TR, and the first 15 bases of the Sp as the primary predictor of the mutagenesis score in this dataset, a feature denoted as $\Delta E_{Avd+TR+Sp} = E_{Avd+TR+Sp} - E_{TR}$.
